# Children’s Hospital Los Angeles COVID-19 Analysis Research Database (CARD) - A Resource for Rapid SARS-CoV-2 Genome Identification Using Interactive Online Phylogenetic Tools

**DOI:** 10.1101/2020.05.11.089763

**Authors:** Lishuang Shen, Dennis Maglinte, Dejerianne Ostrow, Utsav Pandey, Moiz Bootwalla, Alex Ryutov, Ananthanarayanan Govindarajan, David Ruble, Jennifer Han, Timothy J. Triche, Jennifer Dien Bard, Jaclyn A. Biegel, Alexander R. Judkins, Xiaowu Gai

**Affiliations:** Department of Pathology and Laboratory Medicine, Children’s Hospital Los Angeles, Los Angeles, CA; Department of Pathology, Keck School of Medicine, University of Southern California, Los Angeles, CA

**Keywords:** SARS-CoV-2, Coronavirus Disease 2019, COVID-19, Phylogenetic, Virus Genome Tracking, Haplotype, CHLA COVID-19 Analysis Research Database (CARD)

## Abstract

Effective response to the Coronavirus Disease 2019 (COVID-19) pandemic requires genomic resources and bioinformatics tools for genomic epidemiology and surveillance studies that involve characterizing full-length viral genomes, identifying origins of infections, determining the relatedness of viral infections, performing phylogenetic analyses, and monitoring the continuous evolution of the SARS-CoV-2 viral genomes. The Children’s Hospital, Los Angeles (CHLA) COVID-19 Analysis Research Database (CARD) (https://covid19.cpmbiodev.net/) is a comprehensive genomic resource that provides access to full-length SARS-CoV-2 viral genomes and associated meta-data for over 30,000 (as of May 20, 2020) isolates collected from global sequencing repositories and the sequencing performed at the Center for Personalized Medicine (CPM) at CHLA. Reference phylogenetic trees of global and USA viral isolates were constructed and are periodically updated using selected high quality SARS-CoV-2 genome sequences. These provide the baseline and analytical context for identifying the origin of a viral infection, as well as the relatedness of SARS-CoV-2 genomes of interest. A web-based and interactive Phylogenetic Tree Browser supports flexible tree manipulation and advanced analysis based on keyword search while highlighting time series animation, as well as subtree export for graphical representation or offline exploration. A Virus Genome Tracker accepts complete or partial SARS-CoV-2 genome sequence, compares it against all available sequences in the database (>30,000 at time of writing), detects and annotates the variants, and places the new viral isolate within the global or USA phylogenetic contexts based upon variant profiles and haplotype comparisons, in a few seconds. The generated analysis can potentially aid in genomic surveillance to trace the transmission of any new infection. Using CHLA CARD, we demonstrate the identification of a candidate outbreak point where 13 of 31 CHLA internal isolates may have originated. We also discovered multiple indels of unknown clinical significance in the orf3a gene, and revealed a number of USA-specific variants and haplotypes.

## Introduction

Coronavirus Disease 2019 (COVID-19), caused by the Severe Acute Respiratory Syndrome Coronavirus 2 (SARS-CoV-2), continues to spread rapidly across the globe (Wu et al, 2020; Zhu et al., 2020). SARS-CoV-2 is a +ssRNA virus that belongs to the genus *Betacoronavirus* within the *Coronaviridae* family, which also includes SARS-CoV and MERS-CoV. As of May 2020, at the time of writing, there have been over 5 million confirmed cases globally. Over one third of the global cases are in the United States of America with the U. S. death toll exceeding 95,000. Since the emergence of widely available genomic sequencing, rigorous genomic surveillance and contact tracing (Gire et al., 2014), coupled with extensive testing have come to play critical roles in understanding disease transmission routes, inferring outbreak points, and taking informed measures in containing infections.

The Department of Pathology and Laboratory Medicine at Children’s Hospital Los Angeles (CHLA) launched the Centers for Disease Control and Prevention (CDC) RT-PCR assay for SARS-CoV-2 on March 13^th^, a SARS-CoV-2 whole-genome sequencing research assay on March 26, and an Anti-SARS-CoV-2 Antibody IgG test on April 10^th^, 2020. Testing was conducted on a broad population including patients at CHLA and other institutions within the Los Angeles metropolitan area. To better understand the spread of the COVID-19 pandemic globally, nationally, and locally, we sought to develop the necessary bioinformatics tools and resources to help guide effective infection control and prevention strategies at our hospital and across Los Angeles County.

There are multiple SARS-CoV-2 data repositories. GISAID (https://www.gisaid.org/; Elbe and Buckland-Merrett, 2017; Shu et al., 2017) is the major repository for rapid sharing of COVID-19 data across the globe. GISAID stores SARS-CoV-2 genome sequences and associated clinical, geographical, and epidemiological data. The NCBI Virus (https://www.ncbi.nlm.nih.gov/labs/virus/vssi/#/) and the China National Center for Bioinformation 2019 nCoV Resource (https://bigd.big.ac.cn/ncov/; Zhao et al., 2020) are other major resources. Across these repositories, however, meta-data and naming conventions are not standardized nor are the data consolidated. To overcome this limitation, the CHLA Center for Personalized Medicine (CPM) created the COVID-19 Analysis Research Database (CARD), and has been continuously collecting, consolidating and curating full-length SARS-CoV-2 data from across the world.

While there is a significant lack of COVID-19/SARS-CoV-2 genomic resources and bioinformatics tools in general, NextStrain is an outstanding exception (https://nextstrain.org/ncov; Hadfield J et al 2018). It serves as a great resource for understanding how the COVID-19 pandemic has been spreading globally. NexStrain tools are visually appealing and highly interactive. However, despite its critical role in the epidemiological studies of COVID-19, it lacks the functionality for analyzing user-supplied data nor does it allow for sophisticated interactive tree manipulation required by scientists for in-depth interrogation of likely infection patterns. Therefore, to complement NextStrain and to serve the advanced needs of scientists, we developed a highly interactive phylogenetic analysis platform. The platform allows for flexible tree operations, interactive displays of much larger trees stratified by various types of metadata, and importantly, provides for interrogation of likely origins of user submitted genome sequences, which are useful in genomic surveillance.

The key to rapid genomic surveillance of a particular genome sequence is to precisely place it within the geographically complete and temporally up-to-date phylogenetic contexts. Combined with accurate clinical records and patient contact history, this would aid in identifying the likely source of infection. For this reason, and assuming the viral sequence can be reliably determined using a next-generation sequencing method of choice by the users, we focused on genome comparison, variant calling, variant annotation, and variant profiling against the global virus isolate collection in order to identify the identical, more ancestral, or likely descendent isolates in order to reveal the potential route of transmission. CHLA CARD can provide such an analysis, taking only a few seconds per genome, to generate and return results dynamically embedded in global, USA, and local phylogenetic trees that can be visualized and manipulated through the web interface.

## Results

### System Design and Modules

Our ultimate goal is to develop a centralized genomic resource to provide SARS-CoV-2 full-length viral sequencing data analysis and reporting functions for both internal and external needs. To accomplish these goals, we needed to a) assemble a comprehensive genomic database of full-length SARS-CoV-2 genome sequences collected from COVID-19 samples worldwide; b) pre-compute baseline phylogenetic trees of published high-quality SARS-CoV-2 sequences globally, nationally, and locally; c) build a Web Portal for interactive querying of SARS-CoV-2 sequences and associated meta-data; d) implement online bioinformatics tools for interactive phylogenetic analysis and visualization; e) develop a suite of bioinformatics tools for SARS-CoV-2 genome comparison, variant calling and functional annotation; f) devise phylogenetic analysis tools to infer transmission patterns and to determine the origins of infections; and g) establish an Application Programming Interface (API) that allows programmatic access to results in JSON format, such that this resource can be integrated into other SARS-CoV-2 analytical pipelines seamlessly. These represent components of our overall system design goal and constitute the modules of CHLA CARD.

### Data Preparation

The CHLA internal SARS-CoV-2 sequencing data were generated using the SARS-CoV-2 whole genome sequencing research assay, established by the CHLA Center for Personalized Medicine and the Virology Laboratory (Pandey *et al*., in preparation). The major external resources of SARS-CoV-2 strains, genome sequences, and variants were GISAID, GenBank, CNCB, and NextStrain. Data were downloaded in various formats depending on the exact implementations of these external resources. The retrieved data were post-processed, merged, reformatted, and loaded into a MySQL database. Within this database, a master table is created to store merged and normalized viral meta-data from these resources based upon a template that is compatible with NextStrain’s meta-data table. This master table is continuously updated to ensure CHLA CARD provides the most up-to-date and comprehensive SARS-CoV-2 data to end users. The viral genome sequences and metadata are cross-examined to identify and to remove duplicate entries present in multiple resources. Variant calls were generated either internally based on the genome sequences or downloaded directly from China National Center or Bioinformation (CNCB), and were formatted by incorporating standard conventions, functional annotations, and virus information.

Thus, the compiled reference database consists of a comprehensive set of full-length SARS-CoV-2 genome sequences, their variants relative to the SARS-CoV-2 reference sequence (NC_045512.2), strains, and associated meta-data. Currently (as of 05/20/2020), CHLA CARD contains data from 30,000 SARS-CoV-2 strains, over 30,000 complete genome sequences, and more than 7,000 variants.

Statistical and categorical analyses of these reported isolates and variants reveal some notable differences between the United States (USA) and non-USA isolates across genes, and for different annotation classes (**Figure 1, Table S1**). For missense variants, the number is highest in orf1ab polyprotein, 1,191 variants were seen at least three times in 21,535 Non-USA isolates and 589 variants in 5,915 USA isolates. This is expected because this is a large gene (21,290 base pairs or 71.1% of the virus genome) (**Figure 1**). The shorter surface glycoprotein S gene which is only 3,822 base pairs in length, contains about 1/5 of the missense variants or 252 for non-US and 106 for USA isolates, at a similar variant density as orf1ab. It is noteworthy that envelope protein gene orf3a’s in-frame deletions and frameshift deletions are only identified in isolates from USA (12), Australia (6), UK (5), China (5) and Spain (2) but not in other countries. There are recurrent deletions around base positions 26157-26161, including 7 frameshift deletions and 4 disruptive and 5 conservative inframe deletions; from both USA and non-USA isolates. The mutation and haplotype analysis using CHLA CARD revealed multiple interesting observations already. The missense mutation frequencies are significantly higher than that of synonymous mutations in the same genes. For example, the numbers are 178 vs 81 (2.2x) in N gene, and 252 vs 144 (1.75x) in S gene for the non-USA isolates (**Figure 1**). Comprehensive haplotype profiling identified wide-spread country- and state-specific haplotypes which together involved about 29.2% of US isolates, which points to the effectiveness of existing travel restriction policies and public health measures in controlling the transmission of SARS-CoV-2 (Shen et al., in submission).

**Figure 1.**
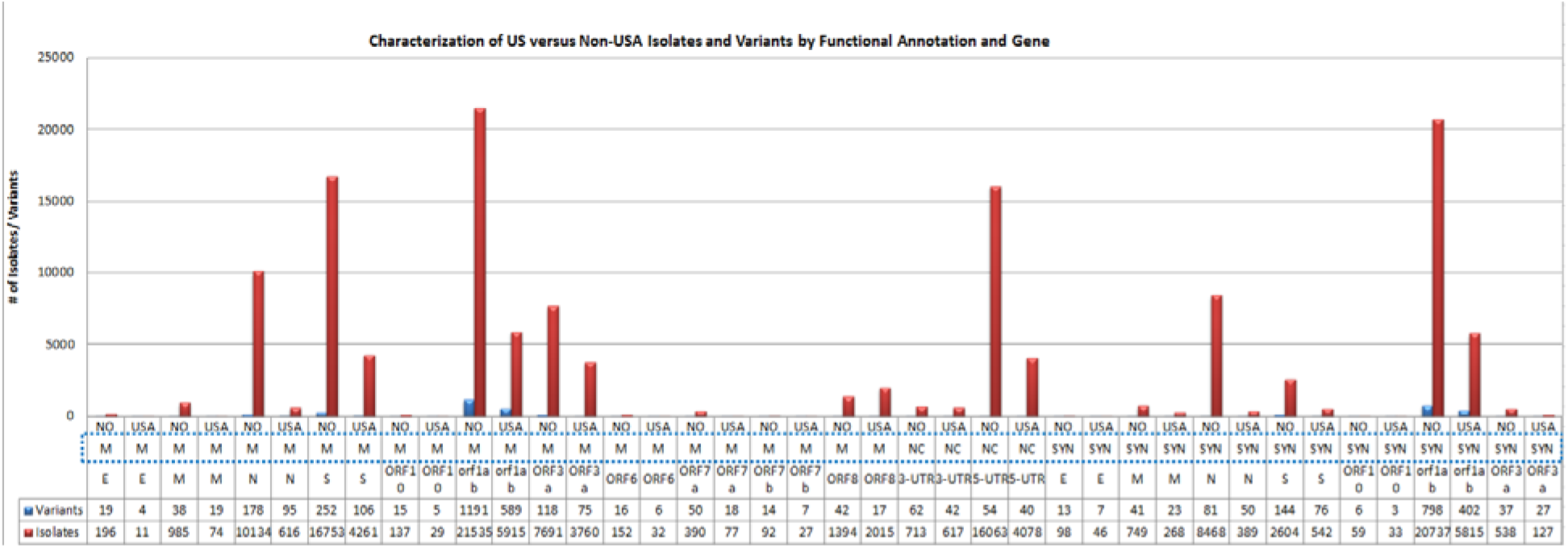
Characterization of SARS-CoV-2 Isolates and Variants by Country, Functional Annotation and Gene. For both USA and non-USA (NO) isolates, variant numbers are highlighted in blue and virus isolate numbers are highlighted in red. Abbreviation: USA = USA isolates, NO= Non-USA isolates, M = Missense, NC = non-coding (intergenic, or orf1ab-5’UTR), FS = frameshift, DEL.IF = inframe deletion, DS = downstream (orf10-3’UTR), SYN = synonymous variants (not shown). Statistics are based on 10,516 high-quality full-genome sequences.

### Phylogenetic Analysis and Reference Phylogenetic Trees

Viral genome sequences are quality benchmarked by counting the missing bases, the length deviation from the reference sequenc (NC_045512.2), and the quality scores (if available). When constructing different subsets of global or country-specific FASTA datasets, the demographic meta-data is used as an inclusion filter for the proper sequences. The reference sequence (NC_045512.2) is always included, and MN996532 (bat/Yunnan/RaTG13/2013) coronavirus sequence is included to serve as the outgroup to root the phylogenetic tree. Multiple sequence alignment (MSA) was conducted with MAFFT version 7.460 (Katoh et al, 2002; Katoh and Toh, 2008) using speed-oriented method FFT-NS-i (iterative refinement method, two cycles) optimized for large datasets. The resulting MSA files were visually examined. All alignments with gaps over 3 bases, or smaller gaps that resulted from inserted “N” (missing) bases, are removed. The 5’- and 3’-end bases are trimmed because they are often of lower quality. The evolutionary history was inferred by using th Maximum Likelihood method and General Time Reversible model (Nei and Kumar, 2000) in MEGA X (Kumar et al., 2018). Initial trees for the heuristic search were obtained automatically by applying Neighbor-Joining and BioNJ algorithms to a matrix of pairwise distances estimated using the Maximum Composite Likelihood (MCL) approach, and then selecting the topology with superior log likelihood value. All positions with over 2% alignment gaps, missing data, and ambiguous bases were excluded. The trees with maximum likelihood were kept as the reference trees for visualization at the web portal. These trees are periodically updated as new genome sequence data are acquired.

### Virus Genome Tracker by Genome Comparison and SNP Profiling against Global Virus Collection

The CHLA CARD Virus Genome Tracker provides: 1) quick comparison of FASTA sequences from complete or partial viral genomes to the reference NC_045512.2 genome for assessing coverage, detecting potential large-scale genome rearrangement, and calling variants; 2) annotation of functional consequence and allele frequency of global virus isolates in order to identify novel and recurrent variants, as well potential candidate variants of clinical significance; 3) identification of strains of identical or similar variant profiles (haplotype) across the country or world, or strains with additional private variant(s) thus likely descendants in the evolutionary tree, or strains with fewer variants but similar haplotype thus likely ancestral in the evolutionary tree. These findings, along with the geographic location, the collection date, and the patient contact history may aid in the tracing of transmission route. The employed strategy is genome comparison followed by SNP profiling, where the query can be the viral sequence in FASTA format, variants in VCF format, or just a free-style list.

The global SARS-CoV-2 isolate collection is matched by shared variant profile of input data. The matched isolates are ranked and classified into 3 match levels: 1) Level 1 matches have identical (Match ratio =1.0), or minimal private variants (Match ratio <1.0) compared with the input isolate and thus could be identical haplotype, 2) Level 2 matches have more additional private variants than Level 1 thus maybe the descendants, and 3) Level 3 matches have fewer variants, and thus may be more ancestral. The Level 1 and 2 top-match isolates are visualized and automatically highlighted on the pre-built reference phylogenetic trees of either Global or USA-specific isolates.

### CHLA COVID-19 Analysis Research Database Web Portal

The CHLA COVID-19 Analysis Research Database (CARD) is implemented as a web service hosted in Amazon Cloud [https://covid19.cpmbiodev.net/]. The back-end database was built using MySQL 5.7 with JSON support. The front-end web interface is written in PHP (www.php.net), JQuery (www.jquery.com), HTML, and JavaScript libraries. Custom bash and Perl scripts are also used at the back-end. The web-based phylogenetic tree visualization and analysis are rendered with Archaeopteryx.js (https://sites.google.com/site/cmzmasek/home/software/archaeopteryx-js) with in-house customization, and taking the prepared phyoXML format file as input (Han and Zmasek, 2009). Genomic visualization is based on the light-weight JBrowse developed by GMOD (http://gmod.org/wiki/JBrowse; Skinner et al., 2009; Buels et al., 2016). Viral genome comparison and variant calling is a wraparound of MUMmer version 4.0.12 (https://mummer4.github.io/; Marçais et al., 2018).

### A Case Study - Phylodynamic Analysis Identifies a Potential Outbreak Point in Los Angeles

The CHLA CPM sequenced 31 viral isolates with its SARS-CoV-2 whole genome sequencing research assay (Pandey et al., in preparation). We analyzed the complete genomes of these isolates by two strategies: 1) Virus Genome Tracker which placed these 31 genomes into the global isolate context in a few minutes, and 2) a refining strategy that merged them onto the reference USA phylogenetic tree, see **Figure 2**. The Virus Genome Tracker strategy using the consensus sequences correctly picked similar external isolates for refined phylogenetic analysis: identical-variant/Level 1 matches (EPI_ISL_415543 & EPI_ISL_415544; **Figure 2**), and descendent/Level 2 match (EPI_ISL_424890 & MT325610). Merging these results onto the reference USA phylogenetic tree we could demonstrate that there is a branching point where 13 of 31 cases center around and make up 9 out of 13 branches from this point. This point is in the middle of the transmission route, inferred from traversing the phylogenetic tree, below a large number of Washington State samples. Among these cases were three members from the same household. This provides a real case example as to how the Virus Genome Tracker and the CHLA CARD facilitated the identification of similar isolates, which can serve as the proxy to query isolates and to place them onto the prebuilt phylogenetic trees in just a few minutes. Customized URLs to the reference trees are automatically generated by Virus Genome Tracker for visualization and exploration.

**Figure 2.**
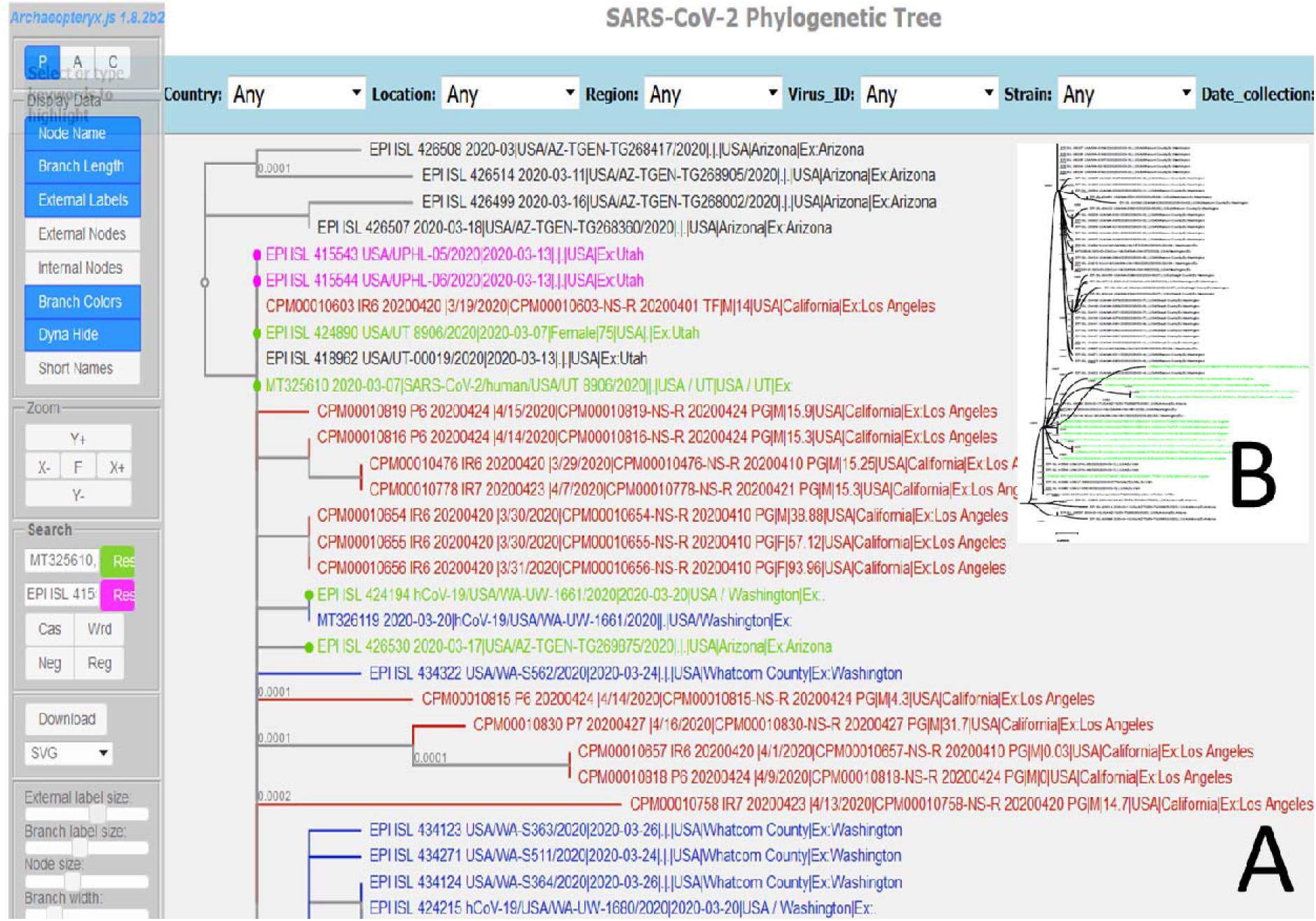
Visualization of USA & Global Phylogenetic Trees by Virus Genome Tracker Showing Queries and Top Matches. **A.** SARS-CoV-2 USA Phylogenetic Tree; **B.** (Inset) SARS-CoV-2 Global Phylogenetic Tree. Internal isolates (green) are mapped against in the phylogenetic context of closest global isolates in (B). Virus Genome Tracker Level 1 and Level 2 matches are automatically highlighted in pink and green, respectively and mapped against phylogenetic context of closest global isolates with query isolates highlighted in red, and other geographically sourced isolates in blue and black in (A).

## Conclusion and Future Plan

CHLA CARD is an integral part of the effort made by the CHLA Department of Pathology and Laboratory Medicine to assemble and to make widely available our strategy and tools for clinical testing, genomic assays, and bioinformatics expertise to address the COVID-19 pandemic. This genomic resource enables and empowers genomic epidemiology studies to understand the origin and spread of SARS-CoV-2 globally, nationally and locally. It also provides in-depth understanding of how the viral genome evolves as the disease spreads across the globe.

The Virus Genome Tracker utilizes MUMmer for super-fast genome alignment and comparison which, combined with the MySQL database backend, can return results within a few seconds (~5 sec) from a search against over 30,000 virus isolates. It supports both whole genome and partial genome sequences, and is thus suitable for handling the consensus sequence data derived from various sequencing strategies. Furthermore, we integrated the genomic variants with gene annotations and population-level data using the smooth scrolling and zooming capability of JBrowse genome browsing for user-friendly browsing and exploration. To our knowledge, there are no published online tools suitable for rapid typing of new SARS-CoV-2 virus isolates, that provide similar functionalities as CHLA CARD, which is empowered by rich viral sequences, isolates, geographic information and phylogenetic data. The Genome Detective Coronavirus Typing Tool (Cleemput et al., 2020) is capable of online sequence but not variant analysis, and has very limited SARS-CoV-2 data with no interactive phylogenetic analysis and visualization.

Together, this genomic resource provides users access to the COVID-19 bioinformatics tools and a comprehensive database of viral genome sequences, including multiple sequence alignments, variants with functional annotations and population frequencies, and precomputed phylogenetic trees, all accessible through the website and the SARS-CoV-2 viral genome browser. Our pilot study using internal cases demonstrates our capability for generating the genomic epidemiology data required to aid in the development of clinical policies and decision making such as devising quarantine and containment strategies.

Modifications to the algorithm and visualization tools employed for virus transmission inference and tracing are in progress to incorporate spatial-temporal-travel data. A mobile version of the web portal is on the roadmap for interactive exploration of data on-the-go. API and beacon capabilities are being implemented to facilitate computational biologists with accessing our standardized viral data.

## Acknowledgements

We are thankful to other members of CHLA’s Virology Laboratory, Division of Laboratory Medicine, and the Center for Personalized Medicine for their strong support of this project. We thank the CHLA Information Security team, including Conrad Band, Adam Miller and William Cox, for their technical support. The phylogenetic tree is rendered with Archaeopteryx.js. The SARS-CoV-2 genomes and meta data were generously shared via GISAID, GenBank, China National Center for Bioinformation (CNCB), Nextstrain and other sources. We gratefully acknowledge the originating and submitting laboratories for making the SARS-COV-2 sequences and associated metadata available via these public resources. For a full list of such contributors, please see from “Virus Isolate” tool on CHLA CARD (https://covid19.cpmbiodev.net/) the full lists of “Originating labs”, “Submitting labs” and “Authors”.

---

**Table S1.** Summary of global SARS-CoV-2 variant collection by gene, annotation and geo-location.

| Gene | Annotation | USA | Variants*** | Isolates |
| --- | --- | --- | --- | --- |
| E | missense | Non-USA | 19 | 196 |
| E | missense | USA | 4 | 11 |
| M | missense | Non-USA | 38 | 985 |
| M | missense | USA | 19 | 74 |
| N | missense | Non-USA | 178 | 10134 |
| N | missense | USA | 95 | 616 |
| S | missense | Non-USA | 252 | 16753 |
| S | missense | USA | 106 | 4261 |
| ORF10 | missense | Non-USA | 15 | 137 |
| ORF10 | missense | USA | 5 | 29 |
| orf1ab | missense | Non-USA | 1191 | 21535 |
| orf1ab | missense | USA | 589 | 5915 |
| ORF3a | missense | Non-USA | 118 | 7691 |
| ORF3a | missense | USA | 75 | 3760 |
| ORF6 | missense | Non-USA | 16 | 152 |
| ORF6 | missense | USA | 6 | 32 |
| ORF7a | missense | Non-USA | 50 | 390 |
| ORF7a | missense | USA | 18 | 77 |
| ORF7b | missense | Non-USA | 14 | 92 |
| ORF7b | missense | USA | 7 | 27 |
| ORF8 | missense | Non-USA | 42 | 1394 |
| ORF8 | missense | USA | 17 | 2015 |
| E | synonymous | Non-USA | 13 | 98 |
| E | synonymous | USA | 7 | 46 |
| M | synonymous | Non-USA | 41 | 749 |
| M | synonymous | USA | 23 | 268 |
| N | synonymous | Non-USA | 81 | 8468 |
| N | synonymous | USA | 50 | 389 |
| S | synonymous | Non-USA | 144 | 2604 |
| S | synonymous | USA | 76 | 542 |
| ORF10 | synonymous | Non-USA | 6 | 59 |
| ORF10 | synonymous | USA | 3 | 33 |
| orf1ab | synonymous | Non-USA | 798 | 20737 |
| orf1ab | synonymous | USA | 402 | 5815 |
| ORF3a | synonymous | Non-USA | 37 | 538 |
| ORF3a | synonymous | USA | 27 | 127 |
| ORF6 | synonymous | Non-USA | 9 | 103 |
| ORF6 | synonymous | USA | 3 | 67 |
| ORF7a | synonymous | Non-USA | 16 | 224 |
| ORF7a | synonymous | USA | 10 | 42 |
| ORF7b | synonymous | Non-USA | 6 | 52 |
| ORF7b | synonymous | USA | 4 | 45 |
| ORF8 | synonymous | Non-USA | 18 | 113 |
| ORF8 | synonymous | USA | 7 | 28 |
| ORF10 | stop_gained; HIGH | Non-USA | 1 | 4 |
| ORF10 | stop_gained; HIGH | USA | 1 | 2 |
| orf1ab | stop_gained; HIGH | Non-USA | 3 | 65 |
| orf1ab | stop_gained; HIGH | USA | 2 | 12 |
| ORF3a | stop_gained; HIGH | Non-USA | 1 | 3 |
| ORF6 | stop_gained; HIGH | Non-USA | 2 | 14 |
| ORF6 | stop_gained; HIGH | USA | 1 | 1 |
| ORF7a | stop_gained; HIGH | Non-USA | 4 | 15 |
| ORF7a | stop_gained; HIGH | USA | 2 | 3 |
| ORF7b | stop_gained; HIGH | Non-USA | 1 | 4 |
| ORF8 | stop_gained; HIGH | Non-USA | 6 | 60 |
| ORF8 | stop_gained; HIGH | USA | 2 | 7 |
| ORF10 | stop_lost&splice_var;<br>HIGH | USA | 1 | 5 |
| E-M | intergenic; MODIFIER | Non-USA | 7 | 104 |
| E-M | intergenic; MODIFIER | USA | 1 | 1 |
| 3-UTR | intergenic; MODIFIER | Non-USA | 62 | 713 |
| 3-UTR | intergenic; MODIFIER | USA | 42 | 617 |
| 5-UTR | intergenic; MODIFIER | Non-USA | 54 | 16063 |
| 5-UTR | intergenic; MODIFIER | USA | 40 | 4078 |
| Coronavirus 3' stem-loop II-like motif (s2m) | intergenic; MODIFIER | Non-USA | 25 | 650 |
| Coronavirus 3' stem-loop II-like motif (s2m) | intergenic; MODIFIER | USA | 12 | 91 |
| M-ORF6 | intergenic; MODIFIER | Non-USA | 2 | 7 |
| M-ORF6 | intergenic; MODIFIER | USA | 1 | 4 |
| N-ORF10 | intergenic; MODIFIER | Non-USA | 12 | 139 |
| N-ORF10 | intergenic; MODIFIER | USA | 7 | 569 |
| ORF3a-E | intergenic; MODIFIER | Non-USA | 4 | 19 |
| ORF3a-E | intergenic; MODIFIER | USA | 4 | 44 |
| ORF6-ORF7a | intergenic; MODIFIER | Non-USA | 2 | 8 |
| ORF6-ORF7a | intergenic; MODIFIER | USA | 1 | 2 |
| ORF7b-ORF8 | intergenic; MODIFIER | Non-USA | 1 | 3 |
| orf1ab | conservative_inframe_deletion | Non-USA | 6 | 97 |
| orf1ab | conservative_inframe_deletion | USA | 5 | 37 |
| ORF3a | conservative_inframe_deletion | Non-USA | 1 | 4 |
| ORF7a | conservative_inframe_deletion | Non-USA | 2 | 8 |
| ORF7b | conservative_inframe_deletion | Non-USA | 1 | 3 |
| orf1ab | conservative_inframe_insertion | Non-USA | 1 | 126 |
| N | disruptive_inframe_deletion | Non-USA | 1 | 4 |
| S | disruptive_inframe_deletion | Non-USA | 3 | 26 |
| S | disruptive_inframe_deletion | USA | 2 | 4 |
| orf1ab | disruptive_inframe_deletion | Non-USA | 10 | 833 |
| orf1ab | disruptive_inframe_deletion | USA | 7 | 23 |
| ORF6 | disruptive_inframe_deletion | Non-USA | 1 | 3 |
| ORF7a | disruptive_inframe_deletion | Non-USA | 1 | 5 |
| ORF8 | disruptive_inframe_deletion | Non-USA | 1 | 8 |
| orf1ab | disruptive_inframe_insertion | Non-USA | 2 | 44 |
| ORF7b | downstream_gene; DISTANCE=32 | Non-USA | 1 | 3 |
| Coronavirus 3' stem-loop II-like motif (s2m) | downstream_gene; DISTANCE=73 | Non-USA | 1 | 3 |
| N | frameshift; HIGH | Non-USA | 1 | 3 |
| S | frameshift; HIGH | Non-USA | 1 | 4 |
| S | frameshift; HIGH | USA | 1 | 1 |
| orf1ab | frameshift; HIGH | Non-USA | 2 | 10 |
| orf1ab | frameshift; HIGH | USA | 1 | 1 |
| ORF3a | frameshift; HIGH | Non-USA | 1 | 3 |
| ORF3a | frameshift; HIGH | USA | 1 | 3 |
| ORF6 | frameshift; HIGH | Non-USA | 3 | 16 |
| ORF7b | frameshift; HIGH | Non-USA | 1 | 3 |
| ORF8 | frameshift; HIGH | Non-USA | 1 | 4 |
| ORF8 | frameshift; HIGH | USA | 1 | 3 |
| ORF3a | initiator_codon; LOW | Non-USA | 1 | 3 |
| CHR_START-orf1ab | intergenic_region;<br>MODIFIER | Non-USA | 2 | 11 |
| ORF10-CHR_END | intergenic_region;<br>MODIFIER | Non-USA | 9 | 54 |
| ORF10-CHR_END | intergenic_region;<br>MODIFIER | USA | 3 | 11 |

\*\*\*: Limite to variants called in three or more isolates

## Appendix A.

## CHLA COVID-19 Analysis Research Database (CARD) Tutorial

## 1. SARS-CoV-2 Virus Genome Browser (Supplemental Figure 1)

1.1. Click the “Virus Genomics” menu, or type the URL: https://covid19.cpmbiodev.net/covid19/cov19snp.php
1.2. Click on any of the “Major Classifications” dropdown. The number of records per category are shown and automatically applied as the filter to limit reportable records as returned by the “2. Keyword Quick Filter”. Composite filter is automatically generated when users select values from multiple drop-down lists.
1.3. Alternatively, click the “Precise Search” button. The search results will contain only those records that exactly match each of the multiple categories from drop-down lists selected at the last step.
1.4. In addition, records can be further filtered by manually typing one or more keywords from the other columns in the search box at “2. Keyword Quick Filter”. Keywords can be partial, and can be a combination of keywords from multiple columns. Add “!” after a word to enable whole word search. Note that the keywords can match any columns shown in the table, whereas the “Precise Search” function matches exactly the same column.
1.5. Save the filtered results by clicking on the “Export Filtered Table to csv” button to save as a csv file to be opened with Microsoft Excel or similar tools.

**Supplemental Figure 1.**
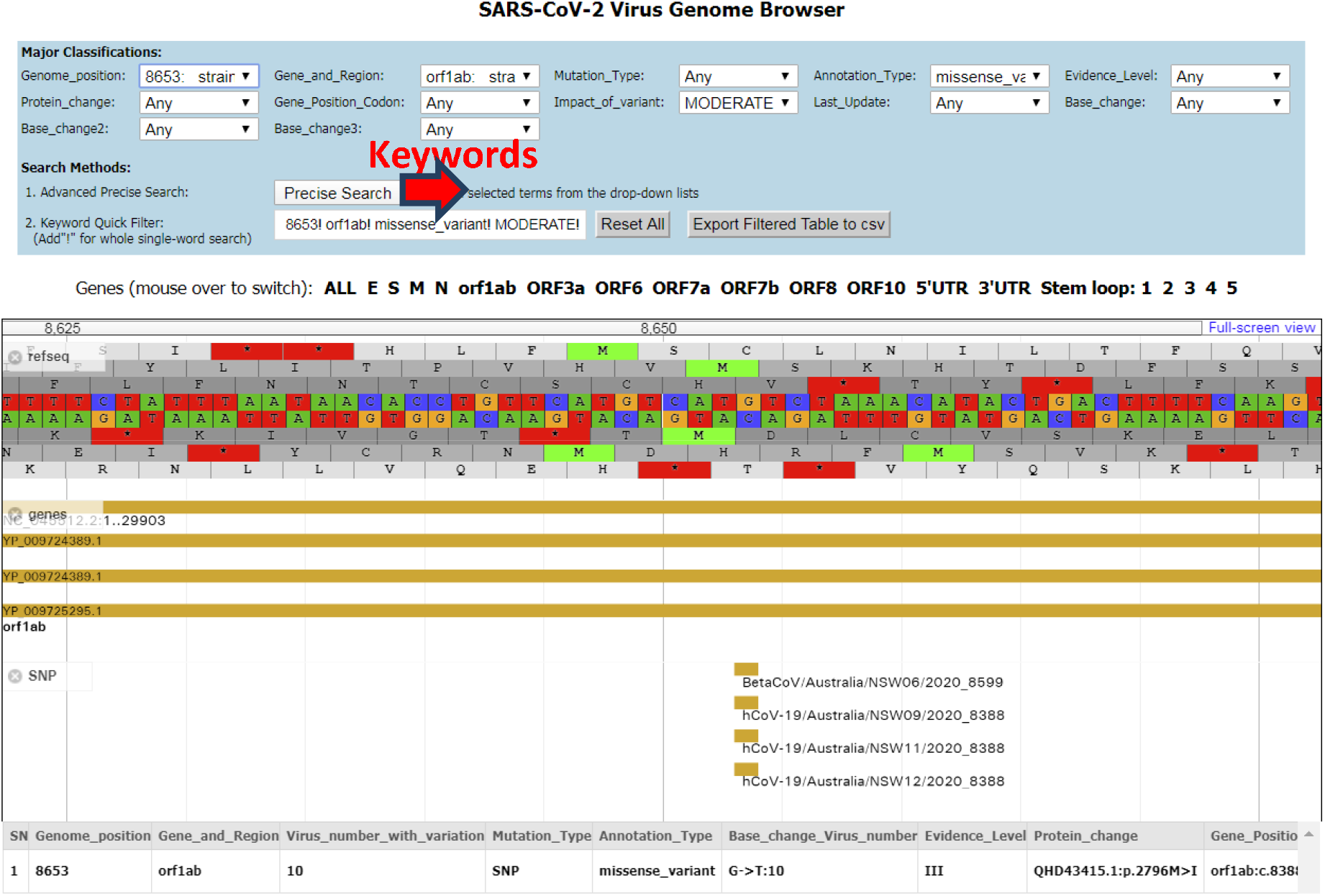
SARS-CoV-2 Virus Genome Browser

## 2. Genomic Visualization in JBrowse (Supplemental Figure 2)

2.1. The genomic data search tool is coupled with a fast JBrowse-based genome browser. The reference RNA sequences, genes, and the variants from global virus collection are shown as tracks by default. The reference RNA sequence is translated in all reading frames for the amino acid being coded. Click on any gene or variant entry to display the full description about the entry in pop-up windows.
2.2. Use mouse to drag-and-move the genomic regions under view by clicking and dragging in the track area or by pressing the left and right arrow keys. Move the mouse to the top of genome browser until a “+” icon appears, then press the mouse to begin dragging to the left or the right to select regions to zoom-in. Users can select smaller region to zoom-in to it up to the base level.
2.3. Changing the variant “Genome_position” or “Gene_and_Region” drop-down list, the genome browser will automatically jump to the specific regions defined by the selected position or gene to display the reference genome and global virus variant reference.
2.4. In addition, all genes and non-coding genomic features are linked above the genome browser. Mousing over these links will switch the visualized region to the gene or “ALL” link for the whole genome view.
2.5. Visualize user’s custom data along with the default reference data provided by the resource.
2.6. Click on the “Full screen View” link at the genome browser’s top-right to go to the full function mode.
2.7. Use the top menu “Track”, select “Open Track File or URL” to upload or link to your custom data. It supports GFF3, BED, FASTA, BAM, and VCF (tabix indexed) formats. Quantitative tracks can be in BigWig and Wiggle formats. Data in BAM, BigWig, and VCF formats are displayed directly from the compressed binary file with no conversion needed.
2.8. For other functions of track manipulation, highlight, resize, keyword and position based search, refer to the “Help” menu for detailed documentation.
2.9. Note that the custom track data is only visible to the user.

**Supplemental Figure 2.**
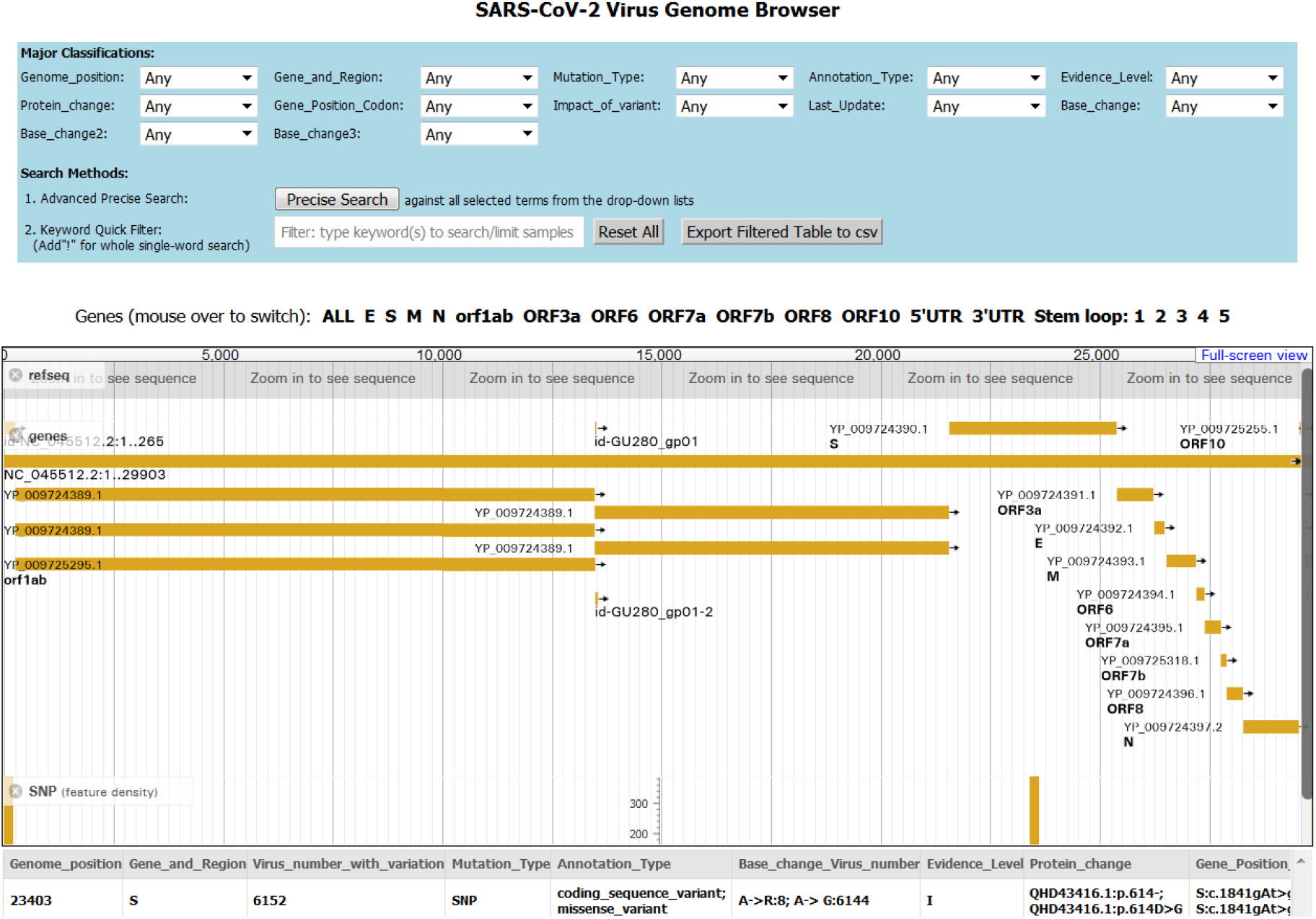
SARS-CoV-2 Virus Genome Browser

## 3. SARS-CoV-2 Virus Isolate Browser (Supplemental Figure 3)

3.1. Click the “Virus Isolate” menu, select “Virus Isolate Browser”, or type the URL: https://covid19.cpmbiodev.net/covid19/cov19strain.php
3.2. Click on any of the “Major Classifications” dropdown. The number of records per category are shown and automatically applied as the filter to limit reportable records as returned by the “2. Keyword Quick Filter”.
3.3. Alternatively, click the “Precise Search” button. The search results will contain only those records that match to ALL of the selected categories from only the drop-down lists in the last step.
3.4. In addition, records can be filtered further by manually typing one or more keywords from the other columns in the search box at “2. Keyword Quick Filter”. Keywords can be partial, or can be a combination of keywords from multiple columns. Add “!” after a word to enable whole word search for it. Note that the keywords can match on any columns shown in the table. Click on “Export Selected” button to save the filtered records as csv file to be opened with Microsoft Excel or similar tools.

**Supplemental Figure 3.**
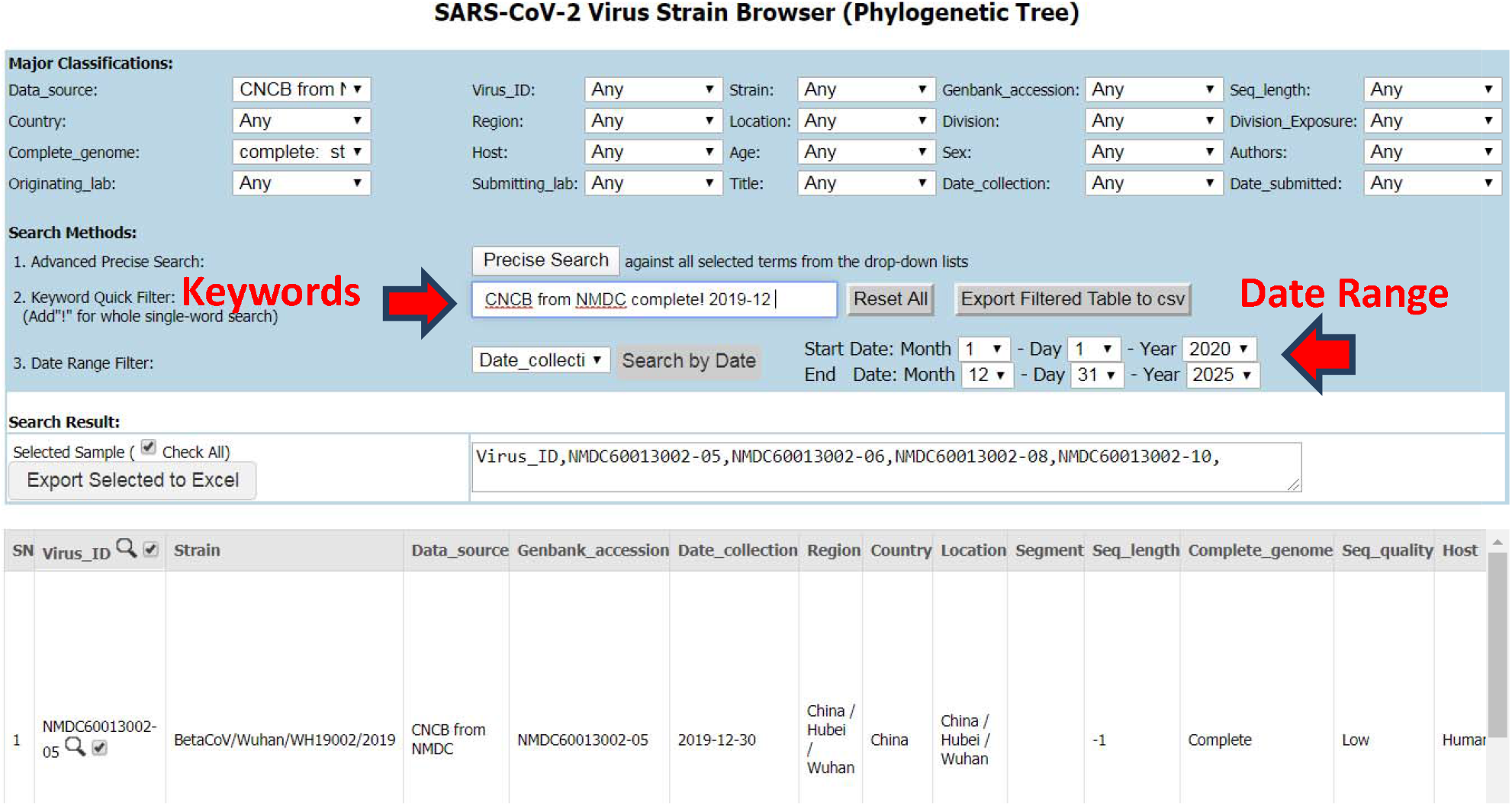
Virus Isolate Browser

## 4. Virus Genome Tracker - Tracking a SARS-CoV-2 Strain by Genome Comparison and SNP Profiling Against Global SARS-CoV-2 Collection (Supplemental Figure 4)

4.1. Type or copy-paste the input to the top text area. The input can be 1) FASTA-format sequence data with header, or 2) VCF-format variant list with or without header, or 3) free style format which is a comma or white space separated variant list where each variant is named as POSITION-REF_Allele-VARIANT_Allele (i.e. 28851-G-T).
4.2. For VCF input, the positions must be based on the reference sequence NC_045512, but the first chromosome column naming is not required to be same as NC_045512. For optimal accuracy in strain matching, variant list from complete genome or high coverage virus genome is preferred.
4.3. Click on the “Annotate” button, check the results of analysis
4.4. For FASTA input, the first part of result is “Genome alignment report” and “SNPs called from your fasta”. The Genome alignment report lists the sequence statistics, comparative genome summary such as coverage on reference and query sequences, chromosomal rearrangements, and SNPs/INDELs called, plus the original sequence. The table of SNPs called lists their positons on both reference and query sequences.
4.5. For all of the input formats, variants are annotated by gene, functional consequence, impact, and global virus isolate allele frequency. Co-located variants are also reported from CNCB data.
4.6. Click “POS”/Search icon in annotation table to visualize its region in the Genome Browse at the bottom.
4.7. The global SARS-CoV-2 isolate collection is matched by shared variant profiles of input data. The top match summary table lists virus collection date and geographic information. Click on strain names to view the full report and genome sequences.
4.8. The matched isolates are ranked and classified into 3 match levels: 1) Level 1 matches have identical (Match ratio =1.0), or minimal private variants (Match ratio <1.0) than the input isolate thus could be identical haplotype; 2) Level 2 matches have more additional private variants than Level 1 thus may belong to the descendants; and 3) Level 3 matches have fewer variants, thus may be located at the more ancestral branches.
4.9. Visualize top match virus isolates on the pre-built backbone reference phylogenetic trees for either Global or USA-specific isolates. Click on the links then in the new windows, and follow the pop-up instruction to examine the trees. Level 1 and 2 will be pre-highlighted on the trees by clicking in new page’s tree panel. More details can be found in section “Phylogenetic Tree Visualization”.

**Supplemental Figure 4.**
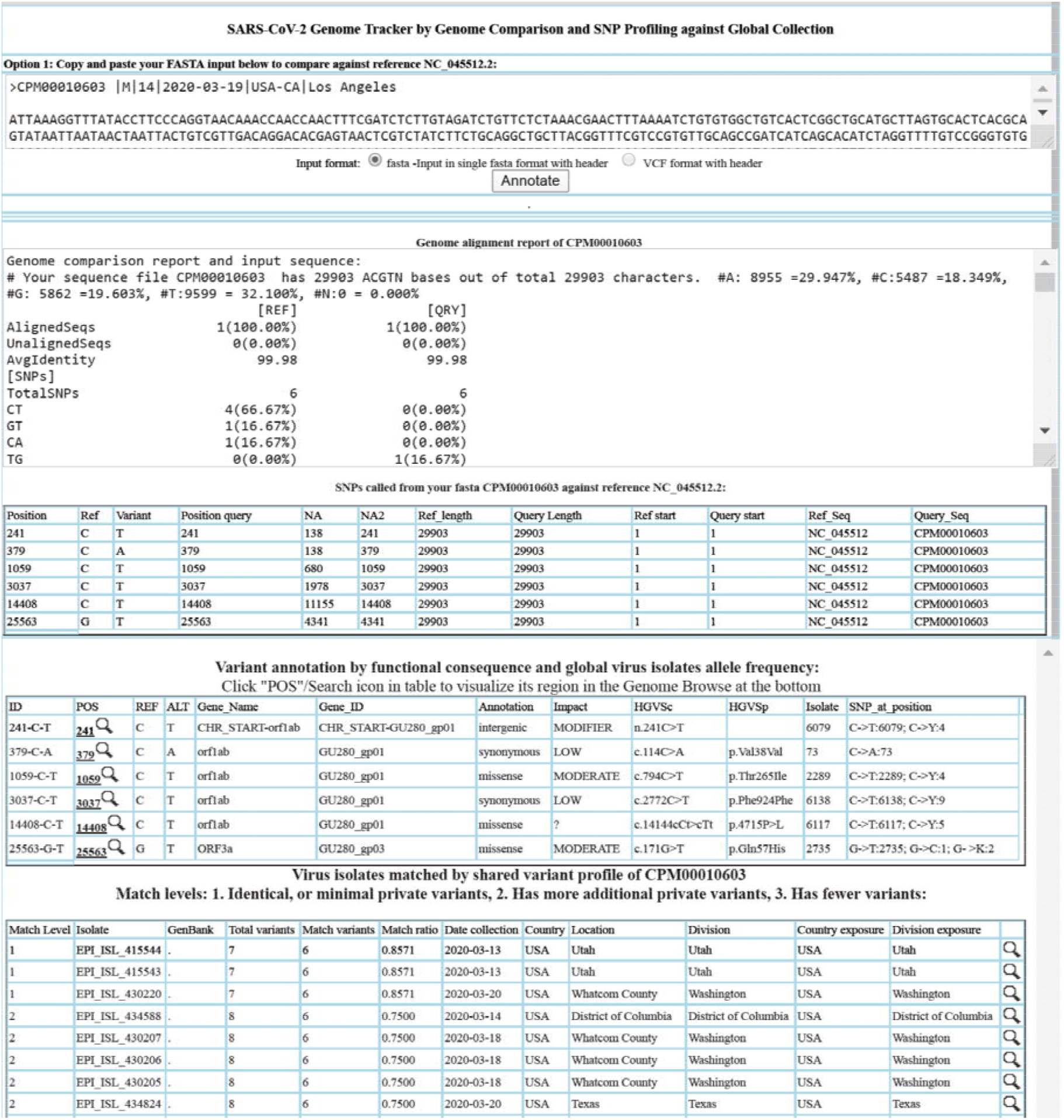
Virus Genome Tracker.

## 5. Online Phylogenetic Tree Visualization (Supplemental Figure 5)

5.1. The tool provides 3 types of phylogenetic trees by default: the regular-mode USA and global trees which are USA-specific isolates, and the global collection. Each of them also has a time series animation mode version.
5.2. Select one of the tree types from the top menu “Phylogenetic Tree” or from the home page, then wait till the tree is fully loaded which could take 30 seconds depending on the network speed, the server usage load, and user’s device performance.
5.3. Virus Genome Tracker initiated visualization starts in the regular tree mode, and it will highlight the Level 1 and Level 2 matches in pink and green respectively and automatically with a click in the tree panel.
5.4. Use drop-down list to search by pre-defined keywords. The current selected keyword and the previous selected keyword will highlight tree nodes in pink and green simultaneously. Available virus meta-data filters include: country, location, region, virus ID, strain name, and date of collection.
5.5. Or manually type keywords to search boxes on the left control panel to highlight strains. If both search boxes at the left control panel have keywords, then matching nodes are highlighted in pink and green simultaneously. For multiple word search: use ‘,’ for logical OR and ‘+’ for logical AND.
5.6. Press mouse in the tree panel and drag to move the graphics. Scroll the mouse middle button to quickly zoom-in and zoom-out the tree.
5.7. Use ZOOM buttons (Y+/− or X+/−) to view more strains, use other left control panel buttons to manipulate the tree/subtree.
5.8. Use the left panel P/A/C buttons to redraw tree to different shapes.
5.9. Click on a tree branch/node/name to check node meta-data, and operate on node/subtree. Sometimes, user needs to use mouse to move the “Node data” popup box away to display the “subtree” operation menu.
5.10. Click on the branch forks to see the number of nodes under the branch. Use the “Go to Subtree” option to quickly check the subtree in zoom-in mode.
5.11. Use the left control panel’s download button to download the current visible tree portion in PNG, or SVG format, or export the subtree as phyloXML and New Hampshire/Newick format to download the tree for offline visualization using own preferred phylogenetic tools.
5.12. In the Time series animation mode, the isolates are colored by date of collection. The temporal sampling starts in December 2019, and increases at an interval of 10 days. The current time point is highlighted in pink, and the previous time point is highlighted in green. The date information is shown in the left panel “Search” boxes. For optimal effect, users may try the left panel P or A buttons to orf3a.
5.13. Redraw tree shapes for better display, then use the ZOOM buttons (X+/− or Y+/−) to adjust width/height. Viewing in zoom-in subtree mode also helps for some clades.
5.14. Multiple highlighting with drop-down list filters

5.14.1. Select 2 filters from the drop-down list at top. They can be from the same list, or from a different list.
5.14.2. Tree is highlighted in 2 different colors simultaneously. The current selection is highlighted pink, and the previous selection is in green.
5.14.3. Type in the left search boxes to revise the filters. Separate the keyword terms with “+” for logical “AND”, or “,” for logical “OR”.

**Supplemental Figure 5.**
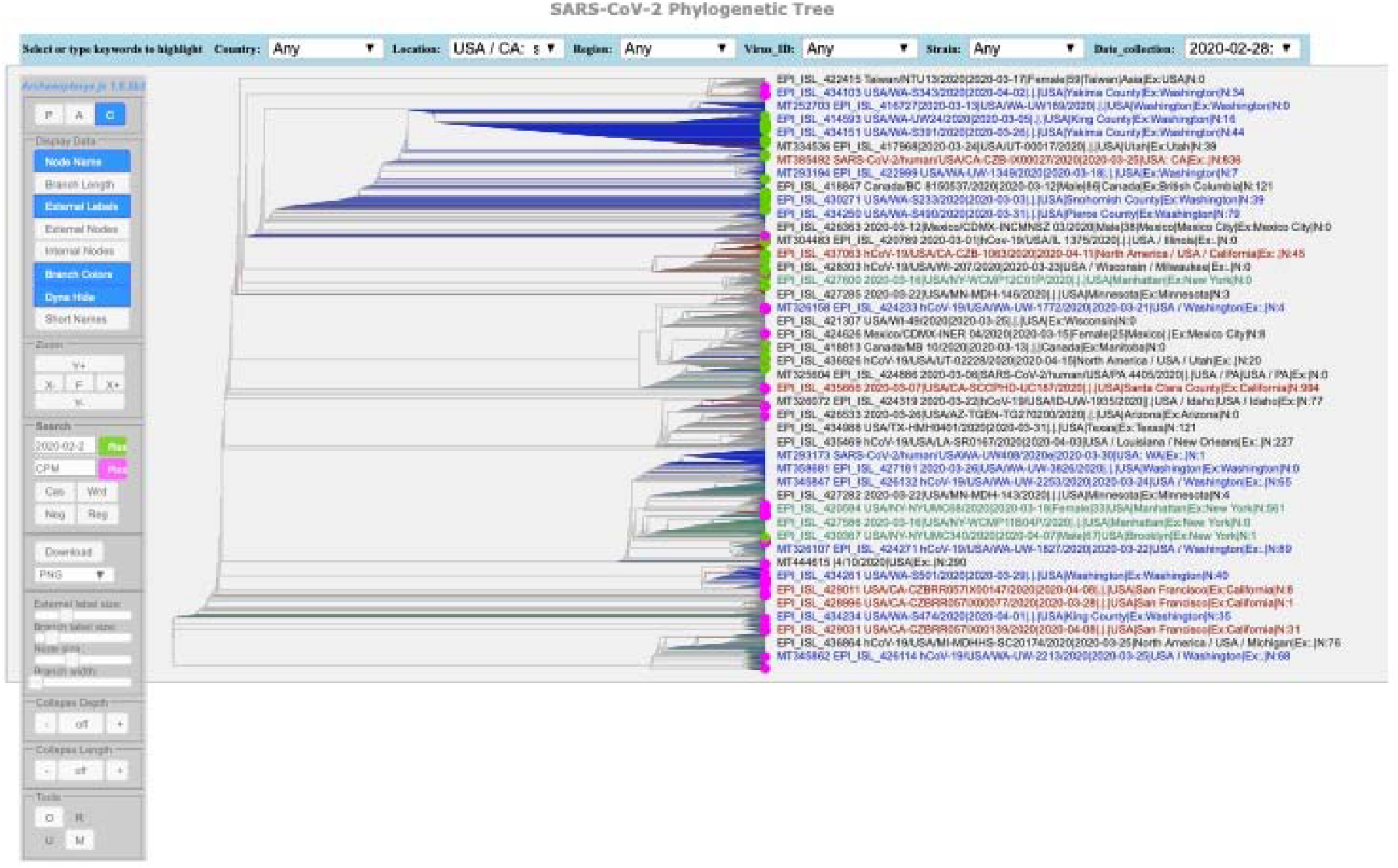
Online Phylogenetic Tree Visualization & Manipulation. The drop-down menus at the top can be used to select and to define two filters for highlighting the tree in different colors simultaneously. The left control panel defines tree manipulation and download options. Further node and subtree manipulation options pop up and become available upon clicking in the tree or on a node.

## Reference

Buels R, Yao E, Diesh CM, Hayes RD, Munoz-Torres M, Helt G, Goodstein DM, Elsik CG, Lewis SE, Stein L, Holmes IH. (2016). JBrowse: a dynamic web platform for genome visualization and analysis. Genome Biol. 17:66. doi:10.1186/s13059-016-0924-1. PMID: 27072794.

Cleemput S, Dumon W, Fonseca V, et al. (2020) Genome Detective Coronavirus Typing Tool for rapid identification and characterization of novel coronavirus genomes. Bioinformatics. 2020, btaa145. doi:10.1093/bioinformatics/btaa145.

Gire SK, Goba A, Andersen KG, Sealfon RS, Park DJ, Kanneh L, Jalloh S, Momoh M, Fullah M, Dudas G, Wohl S, Moses LM, Yozwiak NL, Winnicki S, Matranga CB, Malboeuf CM, Qu J, Gladden AD, Schaffner SF, Yang X, … Sabeti PC. (2014). Genomic surveillance elucidates Ebola virus origin and transmission during the 2014 outbreak. Science (New York, N.Y.), 345(6202), 1369–1372. https://doi.org/10.1126/science.1259657.

Hadfield J, Megill C, Bell SM, Huddleston J, Potter B, Callender C, Sagulenko P, Bedford T & Neher RA. (2018). Nextstrain: real-time tracking of pathogen evolution. Bioinformatics (Oxford, England), 34(23), 4121–4123. https://doi.org/10.1093/bioinformatics/bty407

Han MV and Zmasek CM. (2009). phyloXML: XML for evolutionary biology and comparative genomics. BMC Bioinformatics 10, 356. https://doi.org/10.1186/1471-2105-10-356. https://bmcbioinformatics.biomedcentral.com/articles/10.1186/1471-2105-10-356#citeas

Katoh K, Misawa K, Kuma K, Miyata T. (2002). MAFFT: a novel method for rapid multiple sequence alignment based on fast Fourier transform. Nucleic Acids Res. 30:3059–3066.

Katoh K and Toh H. (2008). Recent developments in the MAFFT multiple sequence alignment program. Brief Bioinform. 9:286–298.

Kumar S, Stecher G, Li M, Knyaz C, and Tamura K. (2018). MEGA X: Molecular Evolutionary Genetics Analysis across computing platforms. Molecular Biology and Evolution 35:1547–1549.

Marçais G, Delcher AL, Phillippy AM, Coston R, Salzberg SL, Zimin A. (2018). MUMmer4: A fast and versatile genome alignment system. PLoS computational biology. 14(1): e1005944. https://mummer4.github.io/.

Nei M and Kumar S. (2000). Molecular Evolution and Phylogenetics. Oxford University Press, New York.

Sagulenko P, Puller V, Neher RA. (2018). TreeTime: Maximum-likelihood phylodynamic analysis. Virus evolution, 4(1), vex042. https://doi.org/10.1093/ve/vex042

Shu Y, McCauley J. (2017) GISAID: Global initiative on sharing all influenza data - from vision to reality. Euro Surveill. 22(13). pii: 30494. doi: 10.2807/1560-7917.ES.2017.22.13.30494. PMID: 28382917; PMCID: PMC5388101.

Skinner ME, Uzilov AV, Stein LD, Mungall CJ, Holmes IH. (2009). JBrowse: A next-generation genome browser. Genome Res. 19(9): 1630–1638.

Wu F, Zhao S, Yu B, Chen YM, Wang W, Song ZG, Hu Y, Tao ZW, Tian JH, Pei YY, Yuan ML, Zhang YL, Dai FH, Liu Y, Wang QM, Zheng JJ, Xu L, Holmes EC, Zhang YZ. (2020). A new coronavirus associated with human respiratory disease in China. Nature. 579(7798):265–269. doi: 10.1038/s41586-020-2008-3. PMID: 32015508; PMCID: PMC7094943.

Zhao WM, Song SH, Chen ML, Zou D, Ma LN, Ma YK, Li RJ, Hao LL, Li CP, Tian DM, Tang BX, Wang YQ, Zhu JW, Chen HX, Zhang Z, Xue YB, Bao YM. (2020). The 2019 novel coronavirus resource. Yi Chuan. 42(2):212–221. doi:10.16288/j.yczz.20-030. PMID: 32102777.

Zhu N, Zhang D, Wang W, Li X, Yang B, Song J, Zhao X, Huang B, Shi W, Lu R, Niu P, Zhan F, Ma X, Wang D, Xu W, Wu G, Gao GF, Tan W; China Novel Coronavirus Investigating and Research Team. A (2020). Novel Coronavirus from Patients with Pneumonia in China, 2019. N Engl J Med. 382(8):727–733. doi:10.1056/NEJMoa2001017. PMID: 31978945; PMCID: PMC7092803.

